# Acute induction of IFNα is responsible for the attenuation of the live measles vaccine

**DOI:** 10.1101/2025.06.09.658652

**Authors:** Jerome M. Edwards, Steve Yoon, Jacqueline Brockhurst, Linda J. Rennick, W. Paul Duprex, Isabelle Coppens, Andrew Pekosz, Diane E. Griffin

## Abstract

The live-attenuated measles vaccine (LAMV) protects against measles in a safe, efficacious and durable manner. However, the mechanism by which LAMV is attenuated is not well understood, contributing to the increased vaccine hesitancy that has caused a worldwide resurgence of measles. Here, we provide a molecular model of attenuation that implicates increased host innate immune responses and increased LAMV susceptibility to those responses. Our study leverages both *in vitro* and *in vivo* models of infection to find that acute induction of innate immunity in immune cells is the critical determinant of LAMV attenuation. We show that LAMV, in contrast to pathogenic wild type measles virus (MeV), causes a strong IFNα response in primary human peripheral blood mononuclear cells, which restricts viral replication and spread in a post-entry manner. LAMV can enter, transcribe viral mRNA and translate viral proteins in immune cells, but is not able to form infectious particles and spread from cell to cell. Specifically, we find that mutations conserved across LAMV strains in the P/V/C and H genes are responsible for the induction of IFNα and the restriction of infectious virus production. In PBMC cultures and in rhesus macaques, LAMV infection also results in the acute induction of proinflammatory cytokines that likely play a role in the immunogenicity of the vaccine. Overall, our study provides a virological and immunological framework for LAMV attenuation which can be leveraged in repurposing the LAMV backbone and in future vaccine design.

## Introduction

Measles is a re-emerging viral disease and a serious threat to public health systems despite the existence of a safe and protective vaccine. Infection with measles virus (MeV), one of the most contagious infectious diseases known to humans, typically presents with rash, fever and respiratory symptoms – and in extreme cases, infection can lead to fatal viral encephalitis or the deadly neurological sequela subacute sclerosing panencephalitis (SSPE) [1-5]. Additionally, MeV infection has both short- and long-term effects on the immune system, including lymphopenia and immune amnesia, which can lead to life-threatening secondary infections [6, 7]. Recently, reductions in vaccine coverage for measles, both in the United States and abroad, have resulted in a significant increase in measles cases and measles-related deaths [8].

Attenuated strains of MeV are the basis of the highly successful worldwide vaccination strategy against measles. The live-attenuated measles vaccine (LAMV), which continues to provide protection against currently-circulating strains of MeV, was developed in the 1960s by passaging a clinical isolate of the virus in cell culture until it no longer caused disease in humans [9-12]. LAMV is amongst the most effective, efficient and safe vaccines developed to date and is one of the very few vaccines that produces robust, life-long immunity with a single dose [12]. Critically, despite its status as the benchmark for safety and efficacy, we have only a rudimentary understanding of the nature of LAMV attenuation and of the mechanistic links between attenuation and vaccine efficacy. Past studies have explored how mutations in different viral genes may be connected to attenuation, but results have generally been inconclusive [13-17]. It is crucial to characterize the molecular, cellular and immunological factors of attenuation to define the correlates of protective immunity and to inform strategies for repurposing the LAMV as a backbone for novel vaccines and in cancer treatment [18-21]. Additionally, a better understanding of how the vaccines work can help overcome growing vaccine hesitancy.

In this study, we leverage *in vitro* and *in vivo* models of MeV infection to define that the main determinant of LAMV attenuation lies in its acute induction of innate immune responses, which directly restricts it from spreading in immune cells. Conserved mutations in several viral genes contribute to inducing an acute IFNα response that is responsible for inhibiting viral replication and spread in a post-entry manner. This innate immune response, which effectively restricts LAMV from generating infectious virus particles in a cell-type specific manner and thereby restricts systemic spread, might also be responsible for eliciting the long-lasting immune memory to measles that is the basis of the efficacy for LAMV.

## Results

Recent studies have suggested that a main determinant of LAMV restriction is its inability to spread systemically and disseminate in secondary lymphoid tissues, such as peripheral blood and the lymph nodes [22-24]. To explore this further, we set out to characterize the dynamics of MeV and LAMV infection in two primary culture systems that model the main tissues affected by measles infection: respiratory epithelium and mononuclear immune cells. In primary rhesus macaque-derived tracheal epithelial cells (TECs) that were differentiated at an air-liquid interface, LAMV grew to significantly higher infectious titers than MeV by day 7 post-infection (Fig. 1a). In contrast, in human peripheral blood mononuclear cells (huPBMC), LAMV infection did not result in significant infectious virus production in either the supernatant or the cell-associated compartment – as opposed to MeV (Fig. 1b). This restriction of LAMV was not unique to a specific cell type amongst huPBMCs, as we observed the same phenotype when separately infecting monocytes and B cells that were magnetically isolated from huPBMC pools (Supplemental Fig. 1a). To ensure that the lack of infectious virus wasn’t due to the vaccination status of our huPBMC donors, the infections were repeated in PBMCs and lymph node cells isolated from rhesus macaques that were certified immunologically naive to measles and there observed the same strain-specific inhibition of infectious virus titers (Supplemental Fig. 1b). Two currently circulating strains of MeV (B3 and D8 genotypes) were tested in addition to our primary MeV strain (Bilthoven, C2), and they were also able to efficiently replicate in huPBMCs (Supplemental Fig. 1c). Interestingly, infection of various different immortalized cells, including monocyte-like THP-1 cells and B cell-like HSB-2 cells, all supported infectious virus production for both MeV and LAMV, indicating that LAMV restriction is specific to primary immune cells (Supplemental Fig. 1c).

**Figure 1.**
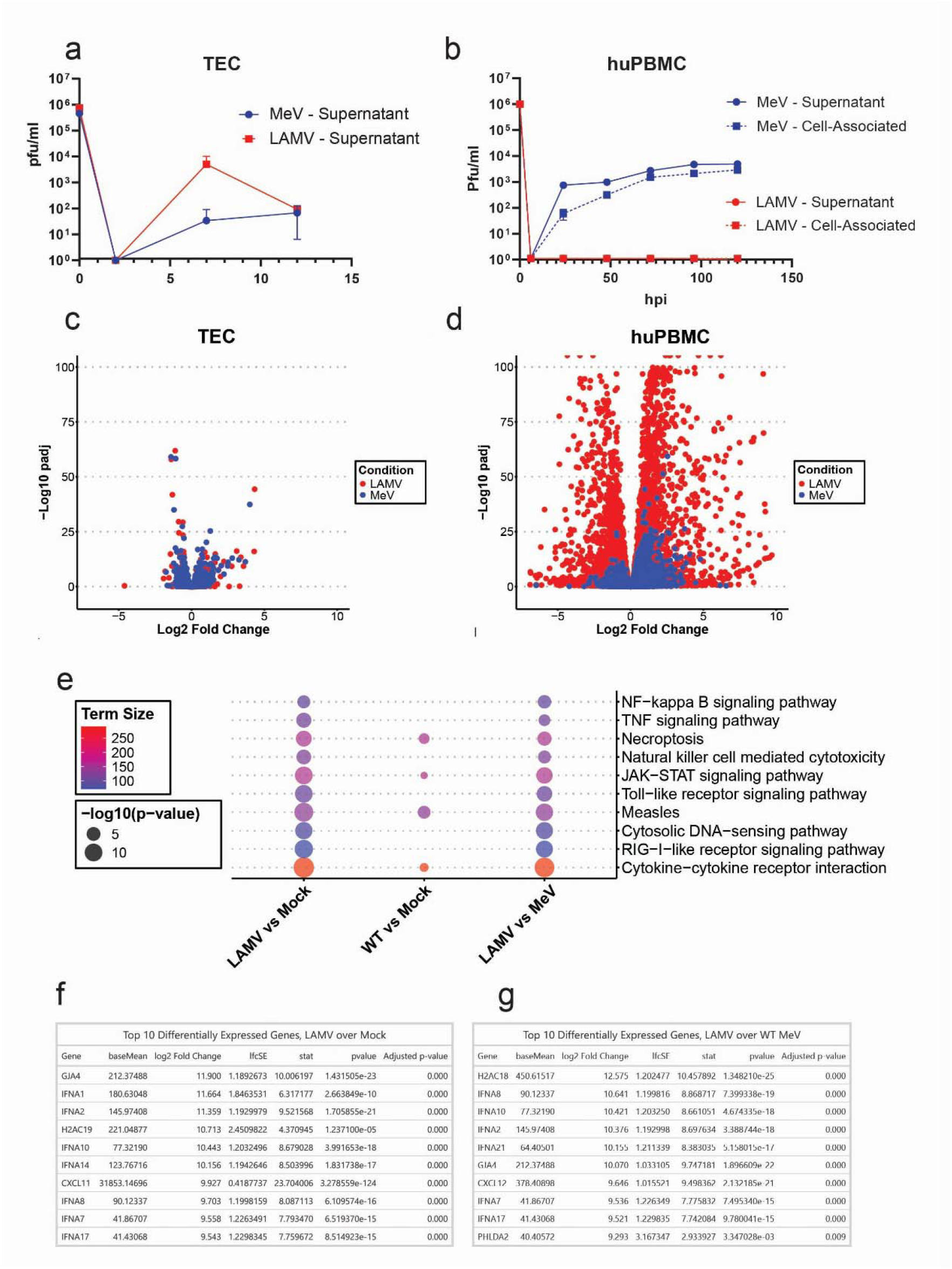
**(a)** Infectious virus titers from the supernatant of primary rhesus macaque-derived tracheal epithelial cells infected basolaterally with MeV or LAMV. **(b)** Infectious virus titers from the supernatant of primary human PBMCs infected with MeV or LAMV. **(c-d)** Volcano plots of differential-gene expression for both MeV and LAMV infection relative to mock-infected TECs **(c)** and huPBMCs **(d). (e)** Gene ontology analysis of enriched pathways for both MeV and LAMV in huPBMCs. Each condition denotes the enrichment of different pathways during infection over another condition. **(f-g)** Tables of top 10 over-expressed genes in huPBMCs during LAMV infection – relative to both uninfected cells **(f)** and MeV-infected ones **(g)**.

To understand the cell type-specific nature of LAMV restriction, bulk RNA-Seq was performed on TECs and huPBMCs at 24 hours after infection with MeV or LAMV. There was only a modest change in gene expression in TECs and no significant differences between cultures that were mock-, LAMV-, or MeV-infected (Fig. 1c, Supplemental Fig. 1g). In contrast, there was a robust transcriptional response in huPBMCs infected with LAMV, relative to both mock and MeV infection (Fig. 1d-e and Supplemental Fig. 1e,f,h). Pathway enrichment analysis showed that LAMV infection of huPBMCs resulted in the significant upregulation of viral-sensing pathways, such as TLR and RIG-I-like signaling cascades, as well as downstream inflammatory pathways such as JAK-STAT and NF-kB (Fig. 1e). More specifically, IFNα subtypes dominated the list of most highly over-expressed genes during LAMV infection, relative to both mock- and MeV-infection, pointing at a type I interferon response potentially being a defining characteristic of LAMV infection in huPBMCs (Fig. 1f-g).

Western blotting was used to determine whether LAMV infection differentially induced the IFN pathway at the protein level (Fig. 2a, Supplemental Fig. 2a-f). By 16 hours, the phosphorylation of TBK-1 and IRF3 was observed during LAMV infection of huPBMCs (Fig. 2a, Supplemental Fig. 2a-b). Robust expression of IRF7 as well as its phosphorylation, were observed in response to LAMV even earlier, at 12 hours post-infection, which in combination with IRF3 is known to induce production of IFNα in cells [25] (Fig. 2a, Supplemental Fig. 2c-d). IFNα and IFNβ levels in the culture supernatant were measured, showing that LAMV infection of huPBMCs, but not MeV, resulted in high levels of secreted IFNα detectable as soon as 1-hour post-infection, and that neither virus significantly induced IFNβ secretion (Fig. 2b, Supplemental Fig. 2g). Consistent with this profile of IFNα expression, STAT1 was phosphorylated very early during LAMV infection (Fig. 2a, Supplemental Fig. 2e). The interferon-stimulated genes OAS-1, RIG-I and MDA-5 were similarly over-induced in LAMV-infected huPBMCs, confirming that interferon signaling occurred downstream of its induction (Supplemental Fig. 2f). Interestingly, MeV infection did not significantly induce the IFN pathway relative to mock-infected cells, which could be attributed to the known immunomodulatory functions of MeV P, V and C proteins [26-28]. To assess the induction of other inflammatory factors, TNFα production was measured, showing an increase during infection with both LAMV and MeV, though the breadth and amplitude of production was greater for LAMV (Fig. 2c). TNFα signaling induces NF-kB-mediated cytokine responses [29]. Consistent with this, a number of antiviral or inflammatory cytokines were released to much greater levels during LAMV infection compared to MeV infection, including factors such as IL-1RA and IL-6, as well as CXCL10, CCL2, 3 and 4 (Fig. 2d, Supplemental Fig. 2h-i). This inflammatory response was also associated with higher levels of cellular cytotoxicity in LAMV infected cultures, as measured by LDH release (Fig. 2e).These data demonstrate that, in huPBMCs, LAMV robustly induces innate immune pathways characterized by a type I IFN axis and the secretion of proinflammatory cytokines.

**Figure 2.**
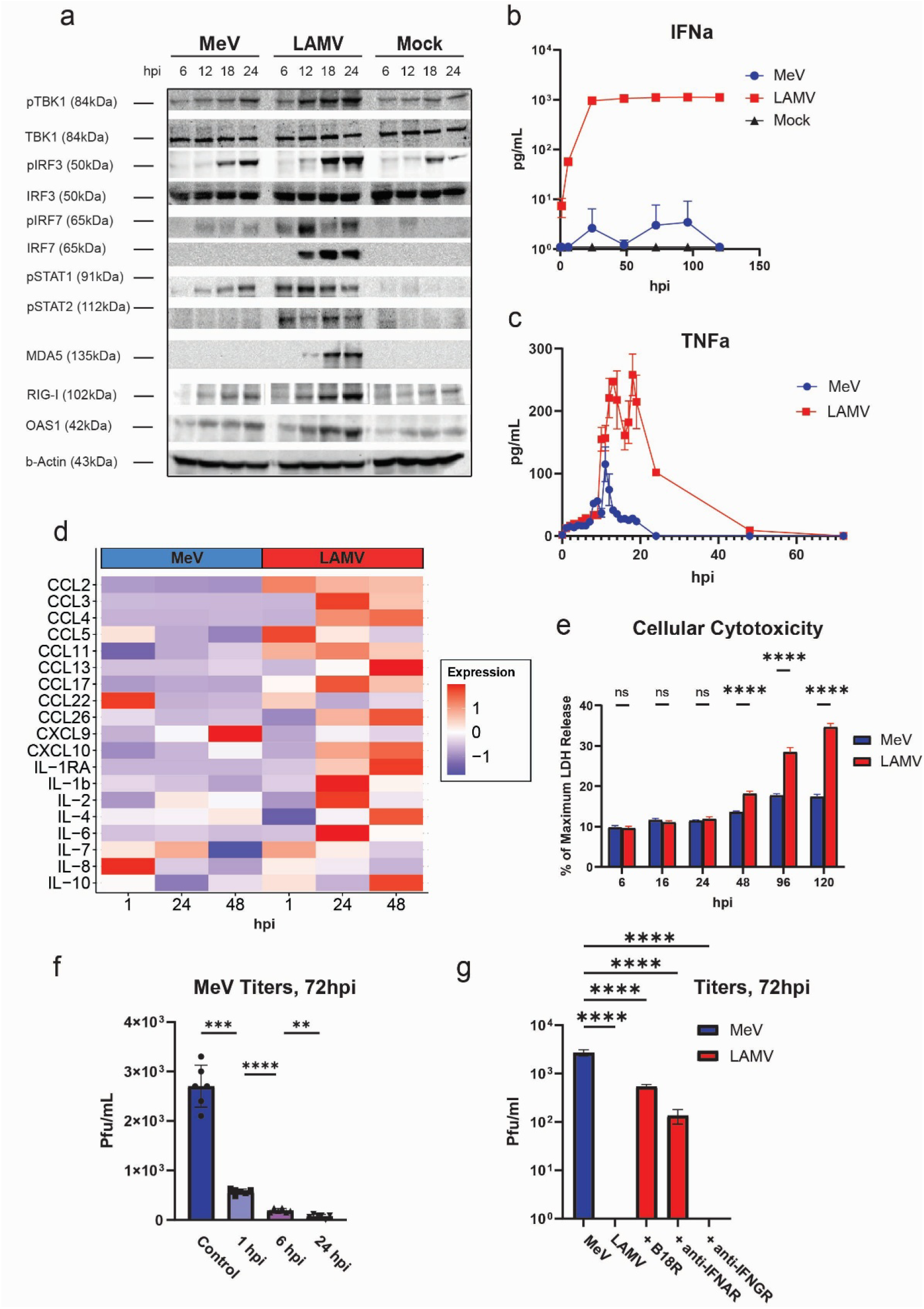
**(a)** Western blot for type I IFN proteins in infected and uninfected huPBMCs at 6, 12, 18, and 24 hours post-infection with MeV, LAMV or mock-infection. **(b)** IFNα secretion during huPBMC infection, by ELISA. **(c)** TNFα secretion during huPBMC infection, by ELISA. **(d)** Heatmap of z-scores from cytokine production, normalized to baseline levels at 1, 24, and 48 hours post-infection of huPBMCs with MeV, LAMV or mock-infection. **(e)** LDH release in huPBMCs, as a percentage of maximum release from cell lysis, during infection with MeV and LAMV. **(f)** Infectious MeV titers, 72 hours post-infection, after resuspension of huPBMCs in inflammatory supernatant from LAMV-infected cells for 1, 6, or 24 hours (MOI=1). **(g)** Infectious virus titers in huPBMCs, 72 hours post-infection, with different anti-IFN treatment.

To determine whether this induction of innate immunity was critical in restricting LAMV infectious virus titers, virus-free culture supernatant from LAMV-infected huPBMCs was collected at 1, 6 and 24 hours and used to treat cultures of MeV-infected cells for 72 hours. The LAMV supernatants from all timepoints were sufficient to significantly reduce MeV titers (Fig. 2f). IFNα was titrated in cells to determine if the amounts of IFNα observed at early timepoints was enough to recapitulate restriction of infectious virus production. Doses of as little as 0.5pg/mL IFNα were enough to restrict MeV infectious virus titers (Supplemental Fig. 2j). Additionally, co-infection of huPBMCs with both MeV and LAMV induced high levels of IFNα and resulted in no infectious virus particles (Supplemental Fig. 2k-l). To confirm that type I IFN was mediating the reduction in infectious virus production, an inhibitory mAb to IFNAR, as well as an IFN decoy receptor (B18R), were used in LAMV infections, with both resulting in a partial rescue of LAMV infectious virus production (Fig. 2g). These experiments indicate that LAMV attenuation is controlled by the early action of IFNα on LAMV-infected cells.

To determine if LAMV was capable of initiating infection in huPBMCs, a strand-specific RTqPCR assay measuring viral gene mRNA was employed. At 6 hours post-infection, expression of 4 mRNA species was detected at equivalent levels in MeV- and LAMV-infected huPBMCs, indicating that LAMV entry and RNA transcription was occurring (Fig. 3a). A time-course of N mRNA expression was then performed in both huPBMCs and Vero cells that express hSLAM – the receptor used by MeV to infect T cells, B cells, dendritic cells, and monocytes [22, 30] - as they are equally permissive to MeV and LAMV infection (Supplemental Fig. 3a). Whereas both viruses accumulate viral mRNA in Vero-hSLAM cells, LAMV specifically was not able to sustain high levels of viral transcription at later timepoints in huPBMCs (Fig. 3b-c). LAMV uses hSLAM in addition to CD46, a complement regulatory protein that is expressed on nearly all nucleated cells, as receptors [31]. To compare both the cell tropism of MeV and LAMV as well as their expression of nucleoprotein N, infected huPBMCs were analyzed by high-parameter flow cytometry, 24 hours post-infection. Overall, 11.36% of total cells were N+ during LAMV infection, as opposed to 3.53% during MeV infection (Fig. 3d). MeV and LAMV nucleoprotein were found in almost all subsets of huPBMCs, including lymphoid and myeloid cells, but LAMV N was consistently present at higher frequencies than MeV N (Fig. 3e, Supplemental Fig. 3b). Additionally, LAMV N was found in cell types that MeV failed to significantly infect, such as NK cells and naïve CD8 T cells (Fig. 3e). This demonstrates that LAMV is able to infect more cell types, and more cells overall than MeV in huPBMC cultures, and that N protein production can occur in LAMV-infected cells. However, whereas more cells were infected by LAMV, the mean MFI of N protein expression was significantly higher for MeV-infected cells across almost all subsets of huPBMCs, suggesting a restriction to the amount of N protein production in LAMV-infected cells (Fig. 3f, Supplemental Fig. 3c).

**Figure 3.**
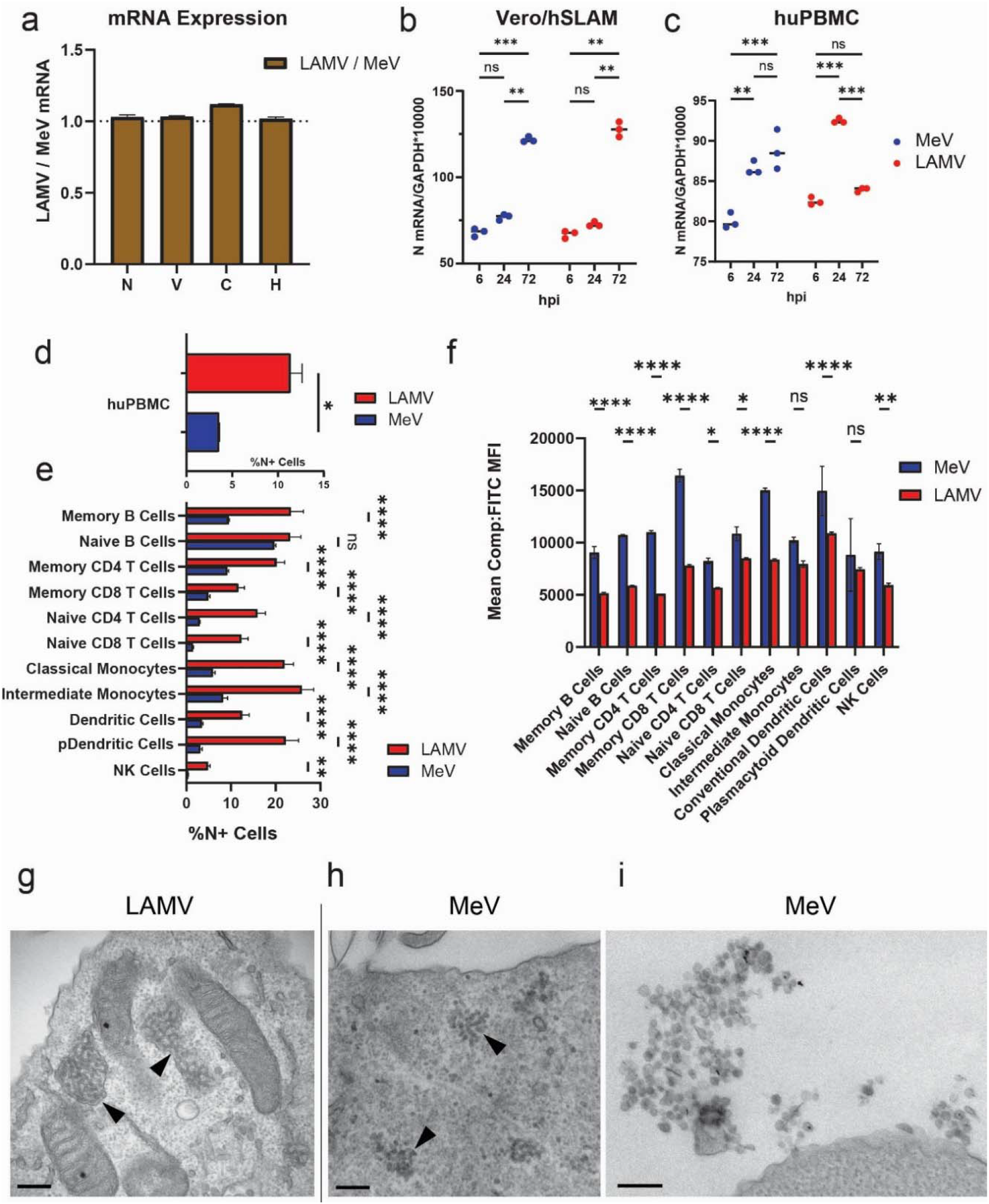
**(a)** Ratio of LAMV to MeV viral mRNA expression for N, V, C and H genes in huPBMCs infected with MeV or LAMV for 6 hours (MOI=1). Viral RNA levels were normalized to GAPDH expression. **(b-c)** Levels of N mRNA in vero-hSLAM **(b)** and huPBMCs **(c)** at 6, 24 and 72 hours post-infection with LAMV or MeV. Viral RNA levels were normalized to GAPDH expression. **(d)** Overall N+ huPBMCs infected with LAMV, MeV or mock-infection for 24 hours, by flow cytometry staining. **(e)** Percentage of N+ cells by different huPBMC subsets during infection with LAMV, MeV or mock-infection for 24 hours. **(f)** Mean compensated FITC MFI for N+ cells in PBMC subsets, where FITC is anti-N **(g-i)** Ultrastructure of LAMV **(g)** or MeV **(h-i)** infection of huPBMCs. Cells were infected for 60h for EM observations, revealing difference in the morphology of intracellular infectious foci (arrowheads) between LAMV and MeV. Egressing viruses were only observed for MeV **(i)**. All scale bars, 0.2 µm.

Ultrastructural studies were conducted to observe differences in intracellular structure and virus particle production between LAMV and MeV in cells. In Vero-hSLAM cells, high amounts of viral RNPs were present in cells infected with both LAMV and MeV (Supplemental Fig. 4a,d). For both viruses, the size and shape of viral RNPs was consistent, and egress events were observable with extracellular virus budding from cells (Supplemental Fig. 4b,c,e,f). In huPBMCs, MeV infection induced RNPs similar in size and shape to those seen in Vero-hSLAM cells (Fig. 3h). Additionally, extracellular RNPs were visible, demonstrating potential egress and infectious particle formation (Fig. 3i). However, LAMV infection resulted in RNP formation, but these were elongated and appeared to be partially membrane-bound in the cytoplasm of cells (Fig. 3g). Importantly, we could not find any evidence of extracellular virus in LAMV-infected huPBMCs, consistent with the lack of infectious virus production. To confirm that this inhibition also restricts viral spread, N-positive huPBMCs were quantified by immunofluorescent microscopy, demonstrating a steady increase in MeV N-expressing cells over the time course of infection, which was not observed in LAMV-infected cells (Supplemental Fig. 3d). These data show that while LAMV can enter, transcribe viral mRNA and produce low amounts of N protein in huPBMCs, the intracellular levels of replication are reduced compared to MeV and contribute to the LAMV attenuation.

To validate LAMV restriction in vivo, two groups of rhesus macaques (n=3 each) were infected intra-tracheally with either MeV or LAMV and followed for two weeks (Fig. 4a). Bronchoalveolar lavage (BAL) samples were collected to understand the dynamics of infection in the respiratory epithelium. Viral RNA was detected in all 3 LAMV-infected macaques but not in MeV-infected animals at one day post-challenge. MeV-infected animals first showed viral RNA at 3 days post-challenge (Fig. 4b). Viral RNA persisted for 14 days in the MeV-infected animals and 2 of three LAMV-infected animals (Fig. 4b). This pattern of viral RNA expression in the BAL was consistent with observations from past macaque studies [23, 32]. However, infectious virus was only recovered from the BAL supernatant of MeV-infected macaques, demonstrating that the vaccine strain is quickly controlled after respiratory infection (Fig. 4c).

**Figure 4.**
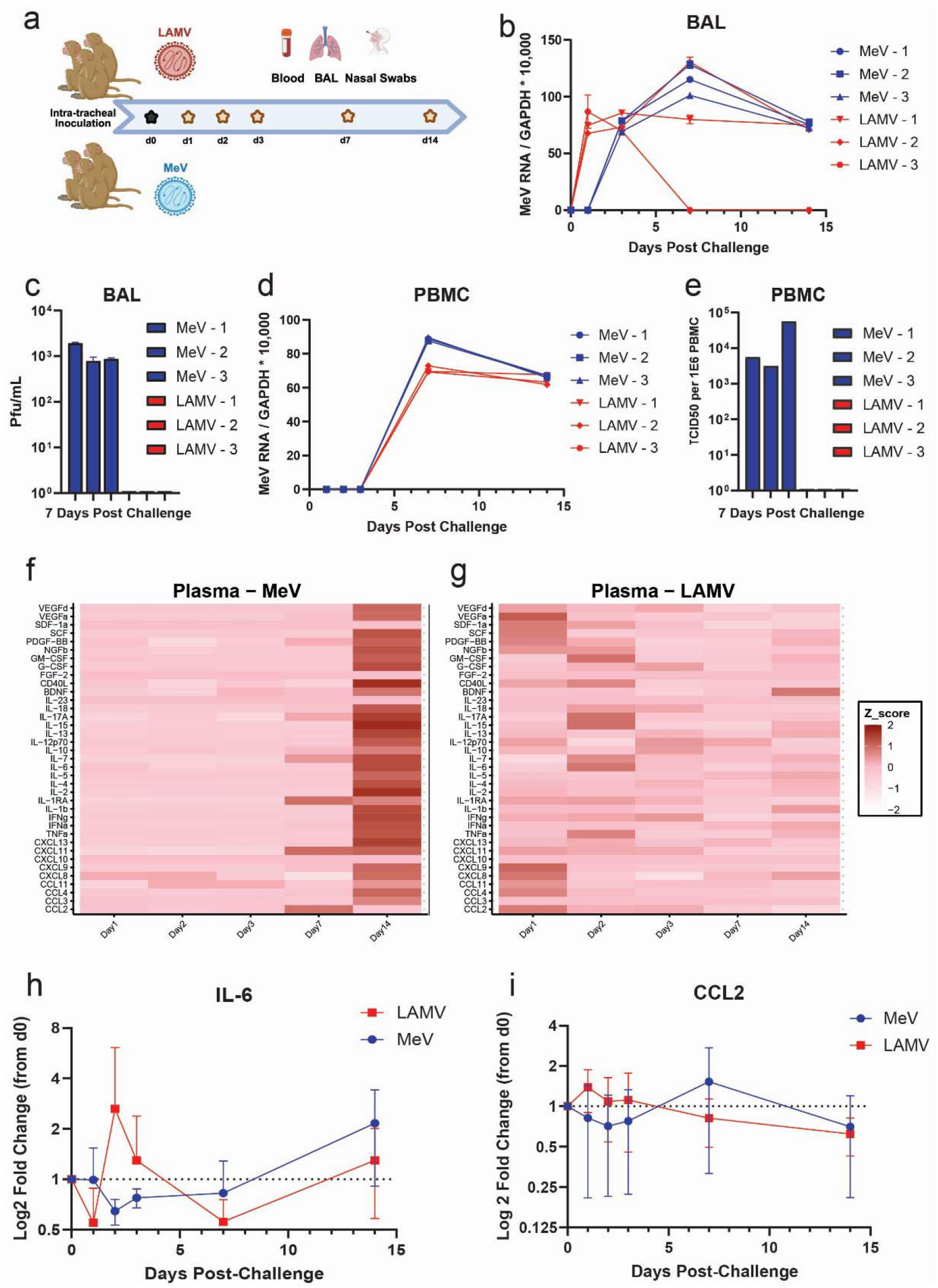
**(a)** Graphical abstract of non-human primate study design. **(b)** RTqPCR of viral RNA in cells collected from bronchoalveolar lavage (BAL) samples for all 6 rhesus macaques. Normalized to GAPDH levels. **(c)** Infectious virus titers in BAL supernatants by plaque assay, at 7 days post-challenge. **(d)** RTqPCR of viral RNA in PBMCs collected from all 6 rhesus macaques. Normalized to GAPDH levels. **(e)** Infectious virus in PBMCs isolated from macaques at 7 days post-challenge, by TCID50. **(f-g)** Heatmap of z-scores of cytokine levels in plasma of both MeV **(f)** and LAMV **(g)** infected macaques, normalized to each macaque’s own baseline expression. **(h-i)** Log2 cytokine expression fold change over baseline (dotted line) for IL-6 **(h)** and CCL2 **(i)**, in MeV or LAMV-infected macaques.

High levels of viral RNA were measured in PBMCs from both groups, starting at day 7 post-challenge, but infectious virus was only detected in MeV-infected macaques (Fig. 4d-e). As shown in *in vitro* infections of huPBMCs, this data is consistent with LAMV infecting PBMCs but being unable to produce infectious virus. All three MeV-infected macaques, but not the LAMV-infected ones, displayed systemic signs of measles infection such as rashes 11 days post-challenge, as well as viral RNA in their nasopharyngeal swabs (Supplemental Fig. 5a-b). This confirmed that disease and systemic spread are specific to MeV, and the lack thereof is a critical determinant of the biological attenuation of LAMV.

To interrogate the innate immunological reactions to both viruses during infection, the expression of various innate and inflammatory factors in the plasma and BAL samples was analyzed. In plasma, MeV-infected macaques developed a strong immune response that encompassed the overexpression of nearly all tested analytes by day 7 or 14 post-challenge (Fig. 4f). This was temporally correlated with the development of viremia and disease. LAMV-infected macaques, who do not develop systemic disease, had an acute response only observable at day 1 and 2 post-challenge (Fig. 4g). This response was characterized by spikes in the expression of pro-inflammatory cytokines and chemokines, such as TNFα, IL-6, CCL2, and CXCL9, which were also induced during *in vitro* infection of huPBMCs with LAMV (Fig. 4h-I, Supplement Fig. 2h-i). In the BAL samples, intermediate levels of inflammatory marker expression were found across the groups and throughout the study (Supplemental Fig. 5c-d). These data demonstrate that the acute inflammation that LAMV induces can be observed in macaques, and that lack of infectious virus production in PBMCs is a critical determinant of LAMV attenuation.

Lastly, the contribution of specific mutations in LAMV to the induction of IFNα and the subsequent restriction in infectious virus production was assessed. Sequence alignments from the different LAMV strains to the sequence of the wild-type Edmonston MeV strain pointed at conserved LAMV mutations in several genes thought to play a role in attenuation (Supplemental Fig. 6a and [15]). A reverse-genetics system was employed to revert these conserved mutations in LAMV back to their MeV amino-acid sequence. Notably, the reversion of LAMV mutations Y481N and G211S in the hemagglutinin, and the G44R, V73A and G225E mutations in the P/V/C genes resulted in significantly reduced levels of secreted IFNα 72-hours post infection of huPBMCs (Fig. 5a). In contrast, reversion of conserved mutations in the matrix gene M (D61G, K89E) resulted in higher IFNα production. The decrease in secreted interferon was correlated with an increase in infectious virus particle production with the H mutations partially rescuing viral titers, and the three mutations in P/V/C fully restoring viral titers by 72-hours post-infection in huPBMCs (Fig. 5b). The mutations in M, as well as all mutations combined, did not rescue infectious virus titers, suggesting that the mutations in M might affect different, synergistic processes that are also essential to viral replication.

**Figure 5.**
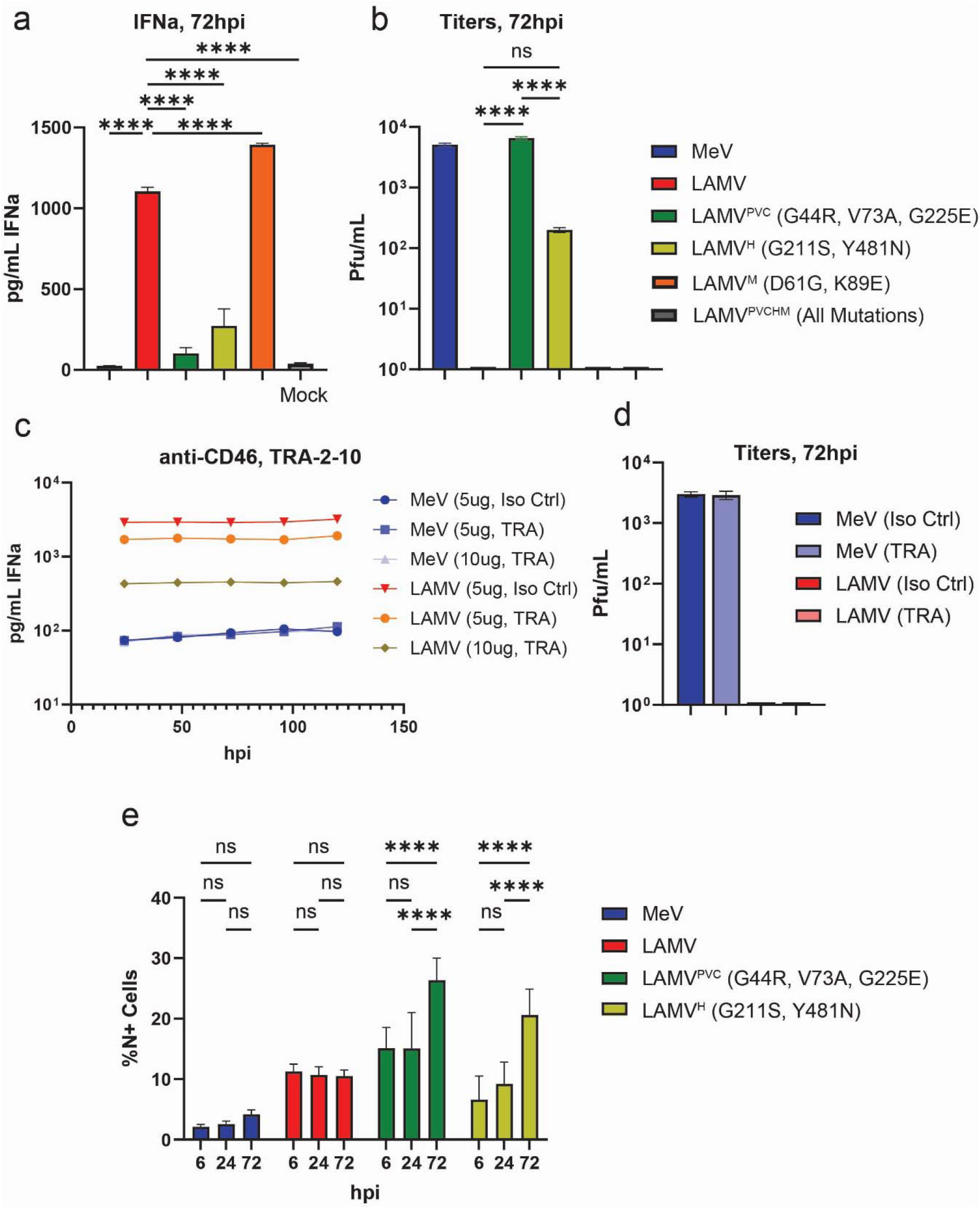
**(a)** IFNα secretion at 72 hours post-infection of huPBMCs with recombinant viruses (MOI=1), by ELISA. **(b)** Infectious virus titers at 72 hours post-infectionof huPBMCs with the recombinant viruses (MOI=1) **(c)** IFNα secretion during the course of PBMC infection with TRA-2-10 antibody against CD46 (or isotype control – ms IgG1κ), by ELISA. **(d)** Infectious titers at 72 hours post-infection during treatment with 10ug TRA-2-10 (or isotype control - ms IgG1κ) **(e)** % N+ cells by immunofluorescent staining at 6, 24, and 72 hours post-infection.

Mutations in the IFN antagonist proteins P/V/C playing a role in altered interferon induction is perhaps obvious: mechanistic changes in their function could quickly result in impaired IFN antagonism or increased sensing of virus infection. That mutations in the viral hemagglutinin contribute to induction of innate immune responses is, however, less intuitive. Because of the Y481N mutation’s known role in CD46 binding, we hypothesized that CD46-mediated-entry might contribute to IFN induction in huPBMCs, as past studies have shown that MeV bound-CD46 can signal through inflammatory pathways and lead to IFN synthesis [33-37]. MeV and LAMV entry through CD46 was inhibited by incubating huPBMCs with monoclonal antibody TRA-2-10, which recognizes the SCR1 domain of CD46 and inhibits H interactions [38, 39]. IFNα induction in the presence of TRA-2-10 was greatly reduced in LAMV but not MeV infection, indicating that H protein interactions with CD46 could trigger IFNα production (Fig. 5c). This did not rescue infectious viral titers (Fig. 5d), as the IFN titrations showed that even small quantities of IFNα can restrict infectious virus production (Supplemental Fig. 2j), but it did confirm that CD46 binding and/or entry by LAMV is at least partially responsible for induction of interferon. Finally, cell entry and spread was quantified for huPBMC infection by recombinant viruses using IF over the course of 72 hours. The mutations in P/V/C resulted in similar initial numbers of N+ cells when compared to LAMV, consistent with the fact that both viruses encode the LAMV H protein (Fig. 5e, Supplemental Fig. 6b). The two mutations in H decreased the initial number of N+ cells to a level closer to that of MeV, which may be attributable to decreased CD46 binding (Fig. 5e, Supplemental Fig. 6b). Unsurprisingly, both sets of mutations allowed for significant levels of viral spread between 24- and 72-hours post-infection, showing that these mutations are indeed critical for LAMV restriction in huPBMCs (Fig. 5e, Supplemental Fig. 6b).

## Discussion

The live-attenuated measles vaccine is amongst the most effective vaccines available to humans as it is safe and confers long-term protection from measles, but the exact nature of its attenuation has not been well characterized. We find that, during infection of both primary human PBMCs and rhesus macaques, LAMV induces the acute production of innate inflammatory factors, which ultimately restricts the systemic spread of the virus. This inflammatory factor induction is characterized by early phosphorylation of IRF7 and IRF3 leading to secretion of IFNα. IFNα restricts viral replication in huPBMCs, though the exact mechanism by which this occurs is still unknown. We show here that the restriction in infectious virus production is post-entry, as LAMV can still enter, express viral mRNAs and translate N protein, but the formation of infectious virus particles is inhibited. Therefore, we find that early IFNα induction is the main host determinant of LAMV attenuation (Fig. 6). That we could not observe this phenotype in immortalized cell lines could explain why past studies in such cells have not shown a strong LAMV attenuation, as transformed cell lines often do not show the same innate immune response to virus infection as primary cells, in large part because the induction and effects of interferons are compromised [14-17, 40, 41].

**Figure 6.**
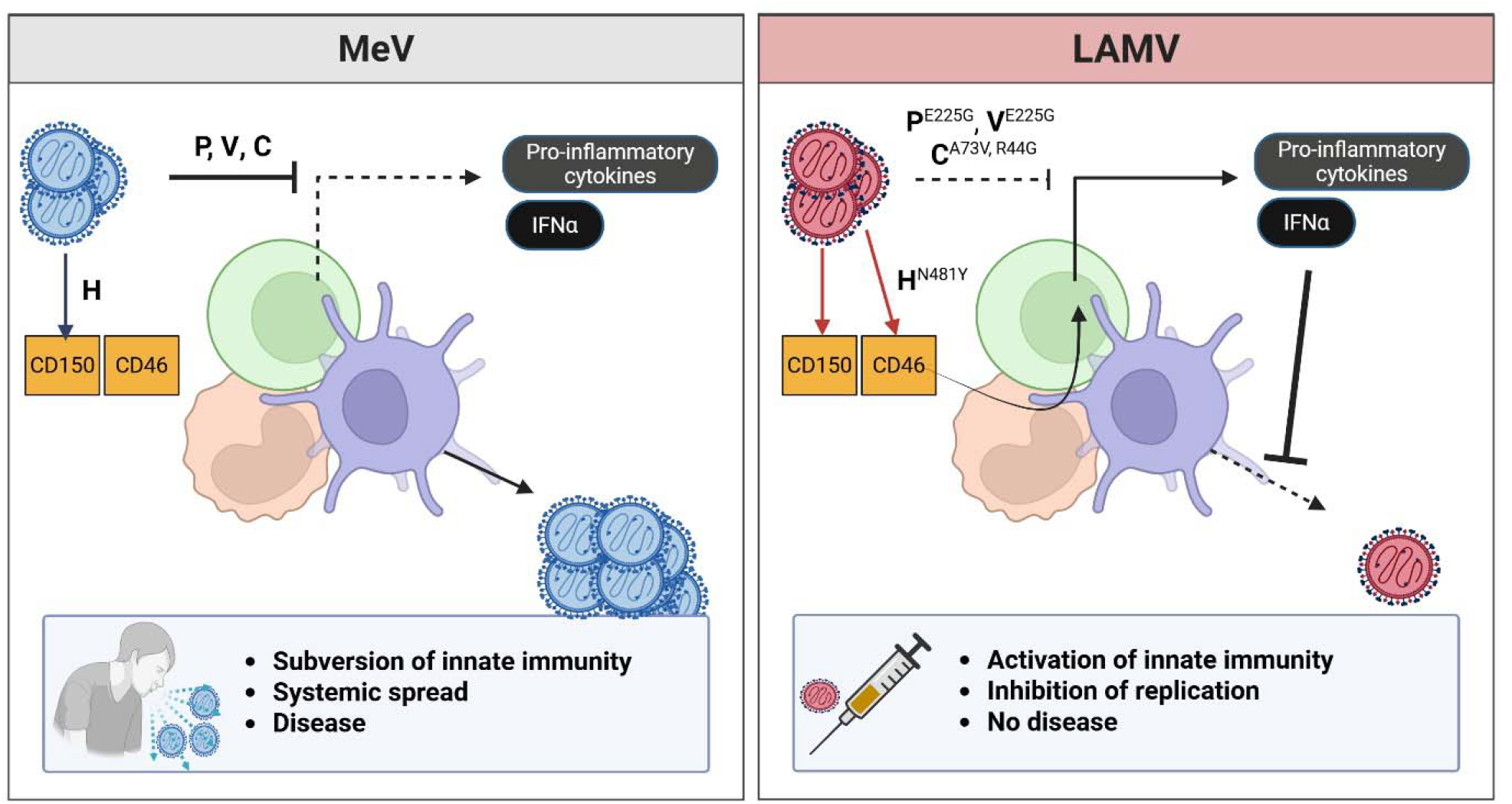
Proposed model of LAMV attenuation relative to MeV infection.

By leveraging conserved mutations in LAMV, we demonstrated that H protein engagement with CD46 as well as P/V/C functions in interferon induction/signaling pathways play a specific role in LAMV attenuation. First, the propensity of attenuated and lab-adapted strains of measles to bind CD46, conferred through several known mutations including Y481N, appears to play at least a partial role in the interferon induction. This is supported by past studies showing that SHP-1 phosphatase, as well as the LCK kinase can be recruited to the cytoplasmic tail of CD46 upon crosslinking to contribute to synthesis of nitrous oxide, IL-12p40 and IFN [33, 35, 36]. Why the G211S/Y481N mutant was able to replicate, though at low levels, and not LAMV during blockage of CD46, might be due to either incomplete inhibition of CD46 entry by TRA-2-10 or other unknown contributions of G211S or Y481N. Additionally, the impact of the P/V/C mutations G44R, V73A and G225E must be further studied. As our phenotype was unique to primary cells, we were not able to efficiently probe functional differences in the proteins, but other studies have already begun to question their potential role in the attenuation of LAMV [42, 43]. These proteins are broad interferon antagonists, whose many targets may offer attractive sites of attenuation. For instance, the G44R mutation has been recently implicated in the knockdown of the C protein nuclear localization signal which is reported to affect its ability to inhibit IFN*β* transcription [43]. Overall, we hypothesize that innate sensing of LAMV, partly achieved through CD46 binding, triggers the interferon signaling pathway which is poorly antagonized by the LAMV P/V/C proteins, allowing it to induce the expression of ISGs that restrict LAMV infectious virus production in immune cells.

Importantly, we also found in our *in vitro* cultures and in the plasma of our macaques that LAMV induces NF-Kb responses, which result in a more systemic inflammation encompassing the release of pro-inflammatory cytokines and chemokines that we hypothesize play an important role in the long-lasting memory responses made to measles during vaccination. However, as these vaccines are given either subcutaneously or intramuscularly, it is important to note that they encounter a different cellular landscape than that of our TEC or huPBMC cultures. We think it possible that they are met by tissue-resident immune cells at the sites of injection, which mount inflammatory responses, recruit leukocytes and control viral spread, altogether resulting in the formation a memory response to the measles vaccines. Overall, the two arms of inflammation mounted against LAMV may be important in their own right: IFN to restrict spread in mononuclear cells, and NF-kB to mount an inflammatory response and improve immunological response at the site of infection.

More generally, these data underline the growing the emergence of innate immunity and interferons as important mediators of attenuation for live-virus vaccines, as it is for the live-attenuated influenza virus (LAIV) and other vaccines [44, 45]. Leveraging the action of type I interferon to limit viral pathogenesis by increasing the induction thereof and/or hampering viral IFN antagonism is an effective strategy for attenuation of infectious virus production, and an attractive option for rational vaccine design. This potential is also underlined by the growing use of IFNs as vaccine adjuvants, given their ability to induce cellular immunity [46]. Though platforms for immunization are diversifying, the ability of live-attenuated viruses to induce robust and long-term immunity at sites of infection without requiring booster doses highlights their enormous potential to limit virus spread and even eliminate infectious diseases such as smallpox [47].

Additionally, recent work shows a correlation between autoantibodies to type I IFNs and adverse reactions to immunization with the 17D live-attenuated vaccine against yellow fever virus [48]. That LAMV is similarly sensitive to innate immune and proinflammatory factors could explain why vaccination can cause disease in immunosuppressed patients such as those with AIDS, whose capacity to mount immune responses is diminished [49, 50]. The implication that alternative vaccination options are necessary for immunocompromised people highlights why it is critical to study the molecular determinants of attenuation for these types of vaccines.

## Supporting information

all supplmental material

## Acknowledgements

We dedicate this paper to Diane Griffin (1940-2024), whose work on the biology of measles infection and alphavirus neurovirulence was truly pioneering and continues to impact us today. We would also like to sincerely thank Debbie Hauer and Haritha Manoj for their lab assistance, Dr. Alan Scott for his guidance and manuscript review, Dr. Elizabeth Thompson for her advice and the use of her flow cytometer, as well as the excellent technical staff (Barbara Smith) of the Electron Microscopy Core Facility at the Johns Hopkins University School of Medicine Microscopy Facility.

## Funding

This work was supported by the National Institute of Health (NIH) grants R01AI153140-05, R01AI182066-02 and T32AI007417-27, as well as by the Merck Company Investigator Studies Program (MISP) grant 102158. The funders of this study had no role in study design, data collection, data analysis, data interpretation, writing of the report, or in the decision to submit for publication.

## Author Contributions

Conceptualization, J.E. and D.E.G.; Methodology, J.E., S.Y., J.B., L.R. and I.C.; Software: J.E.; Validation, J.E., S.Y. and I.C.; Investigation, J.E., A.P. and D.E.G.; Resources, P.D., A.P. and D.E.G.; Writing—Original Draft Preparation, J.E. and A.P.; Visualization, J.E., I.C. and A.P.; Supervision, A.P. and D.E.G.; Project Administration, A.P. and D.E.G.; Funding Acquisition, A.P. and D.E.G. All authors have read and agreed to the published version of the manuscript.

## Competing Interests

D.E.G. was a member of the advisory boards for GlaxoSmithKline, Academia Sinica, and the University of Vermont and has also received funding as a consultant from Gilead, MeVox, Merck, and the US National Institutes of Health. A.P. serves as a consultant for GlaxoSmithkline, Dynavax and Sanofi-Pasteur. All other authors declare no competing interests.

## Data and Materials Availability

Raw FASTQ data for huPBMC and TEC Bulk RNASeq can be found in the Gene Expression Omnibus database under the accession number: (xxx). All scripts used for subsequent analysis are available on GitHub under: (xxx). All source data is included in the supplemental materials and is available under the accession (doi.xxx).

## Supplementary Materials

Materials and Methods

Figures S1-S6

