## Supplementary material for "Acute induction of IFNα is responsible for the attenuation of the live measles vaccine": all supplmental material

**Materials and Methods**

**Animals and Study Design**

Six juvenile (2–3-year-old, 4 males, 2 females) MeV-seronegative, Indian-origin rhesus macaques (*Macaca mulatta*) were obtained from the Johns Hopkins University Primate Breeding Facility. Animals were housed in stable social groups throughout the study. Macaques were sedated for all procedures with ketamine (5-10mg/kg) or ketamine plus dexmedetomidine (0.025 mg/kg) intramuscularly. All studies were performed within an AAALAC-accredited facility in accordance with experimental protocols approved through the Johns Hopkins University Animal Care and Use Committee and guidelines in the Animal Welfare Regulations and *Guide for Care and Use of Laboratory Animals*.

The macaques infected intratracheally with 10e4 PFU of either MeV (Virginia 40.21/2, B3 genotype) or LAMV (Edmonston-Zagreb strain). At selected timepoints, whole blood was collected from the femoral vein into heparin-coated tubes, and plasma was separated by centrifugation. PBMCs and plasma were isolated by whole blood gradient centrifugation on Lympholyte-Mammal (Cedarlane Labs). BAL samples were obtained with a modified catheter by mini-BAL method [1] and nasopharyngeal samples were collected using nylon flocked swabs (Puritan Medical Products) and placed in 1x PBS. Studies were not blinded, and no animals were excluded from analysis.

**Viruses and Cells**

MeV strains Bilthoven (WT, genotype C2 - a gift from Albert Osterhaus, Erasmus University, Rotterdam, Netherlands), M-12 (WT, genotype D8 – a gift from Paul Rota, CDC, USA), Virginia 40.21/2 (WT, genotype B3 – also from Paul Rota) and Edmonston-Zagreb (EZ, LAMV – from the Serum Institute of India) were propagated and assayed by plaque formation in Vero cells stably expressing the human MeV receptor SLAMF1 (Vero/hSLAM cells) [2].Virus used for rhesus macaque infectious was grown in phytohemagglutinin-stimulated human cord blood cells in 1mL PBS. Virus stocks were documented free of defective interfering RNA and mycoplasma. Vero/hSLAM cells and H358 cells (ATCC) were cultured in Dulbecco’s modified Eagle medium (DMEM) with 10% heat-inactivated FBS, 1% L-glutamine, and 1% penicillin-streptomycin. THP-1 cells and HSB-2 cells (ATCC) were maintained in RPMI 1640 supplemented with 10% heat-inactivated FBS, 1% L-glutamine, and 1% penicillin-streptomycin. Peripheral Blood Mononuclear Cells (PBMCs) were isolated from Leukopaks of healthy donors (American Red Cross) by density centrifugation with Lympholyte (Cedarlane) and resuspended in RPMI 1640 supplemented with 10% heat-inactivated FBS, 1% L-glutamine, and 1% penicillin-streptomycin. Isolation of monocytes and B cells was magnetically performed using Dynabeads Untouched Human Monocytes and B cell kits (Thermo Fisher). Experiments were conducted with PBMCs from 3 donors. Cells were infected after a PBS wash by incubation with virus for 1 hour, followed by 2x PBS washes and resuspension in infectious media (IM) consisting of RPMI 1640 or DMEM supplemented with 2% FBS, 1% L-glutamine and 1% penicillin-streptomycin.

The culture and infection of primary rhesus macaque tracheal epithelial cells (TECs) was performed as previously described [3]. Briefly, cells were obtained during necropsy of rhesus macaques, expanded, differentiated for 3-4 weeks at air-liquid-interface and cultured on Transwell inserts. Infections were performed through the basolateral side, using different macaque donors as biological replicates.

**RTqPCR**

MeV RNA was detected and measured in both macaque tissues and infected cells by RT-PCR for the N gene as previously described [4]. Briefly, RNA was extracted, and the N gene was amplified from cells or tissues using TaqMan primers and probe. Data were normalized to the glyceraldehyde-3-phosphate dehydrogenase (GAPDH) control and expressed as (MeV N RNA/GAPDH RNA) × 10000.

For strand-specific RTqPCR of MeV mRNA species, a measles-specific protocol was derived from Kawakami et al [5]. Briefly, reverse transcription of individual viral mRNAs was achieved using tagged primers. Real-time PCR was then performed with the tag sequence as the forward primer and a segment-specific reverse primer to ensure appropriate specificity. All primers are listed in Table 1.

| Target | Purpose | Primer Name | Sequence (5’-3’) | Position (nt) |
| --- | --- | --- | --- | --- |
| N RNA | One-Step RTqPCR | MeV_F | GGGTACCATCCTAGCCCAAATT |  |
| N RNA | One-Step RTqPCR | MeV_R | CGAATCAGCTGCCGTGTCT |  |
| N/A | Real-Time Forward Primer | mRNA_Tag | CCAGATCGTTCGAGTCGT | N/A |
| N mRNA | Reverse Transcription Primer | N_mRNA_Tag | CCAGATCGTTCGAGTCGTTTTTTTTTTTTTTTTTCTAGTCTAG | 1677-1685 |
| V mRNA | Reverse Transcription Primer | V_mRNA_Tag | CCAGATCGTTCGAGTCGTTTTTTTTTTTTTTTTTTTATTCTGGGA | 2697-2705 |
| C mRNA | Reverse Transcription Primer | C_mRNA_Tag | CCAGATCGTTCGAGTCGTTTTTTTTTTTTTTTTTTCAGGAGCTC | 2381-2389 |
| H mRNA | Reverse Transcription Primer | H_mRNA_Tag | CCAGATCGTTCGAGTCGTTTTTTTTTTTTTTTTTCTATCTGC | 9117-9124 |
| N mRNA | Real Time Reverse Primer | N_mRNA_Rev | CTGCAAGCCATGGCAGGAA | 1599-1617 |
| V mRNA | Real Time Reverse Primer | V_mRNA_Rev | GAACCATCAGGGCCAGGT | 2587-2604 |
| C mRNA | Real Time Reverse Primer | C_mRNA_Rev | GAACCTGATACCGAGGGATATG | 2275-2296 |
| H mRNA | Real Time Reverse Primer | H_mRNA_Rev | TGCTTCACATGGGACCAAA | 8978-8996 |

**Supplementary Table 1:** RTqPCR Primer List

**Plaque Assays**

To quantify infectious particles, supernatants and/or cell-associated lysates from infected cells were serially diluted in IM and added to 90% confluent Vero/hSLAM cells in six-well plates and incubated at 37 °C with 5% CO2 for 1 h before overlaying the cells with prewarmed methylcellulose in MEM with 2% FBS, 1% L-glutamine and 1% penicillin-streptomycin. After incubation for 5 days, cells were fixed with 10% formaldehyde, stained with crystal violet and plaques were counted. Amounts of infectious virus are expressed as plaque-forming units (PFU)/mL.

**RNA Sequencing and Analysis**

Total RNA was extracted from pelleted cells and purified from TECs and PBMCs using the RNeasy Mini kit (Quiagen), including on-column DNAse treatment. Quantitation of total RNA was performed with the Qubit BR RNA Assay kit and Qubit Flex Fluorometer (Invitrogen). Library preparation with NEBNext Ultra II directional RNA Library Prep Kit, quality control with Agilent 2100 Bioanalyzer, and sequencing with an Illumina HiSeq machine (2 × 150 nt) were carried out by Admera Health (South Plainfield, NJ, USA).

Fastq files were uploaded to the JHPCE cluster and initial QC on all samples was performed using FastQC. Reads were aligned to the hg38 transcriptome using Salmon 1.10.2 and imported as transcript expression values using tximport [6]. Principle component analysis (PCA) of normalized gene counts was performed using plotPCA and differentially expressed gene (DEG) analysis was performed using DESeq2, during which DEGs were calculated using an alpha of 0.01 and a log2FC cutoff of 1 [7]. Gene set enrichment analysis was performed using gprofiler2 [8].

**LDH Assay**

Culture supernatants were collected and tested for lactate dehydrogenase activity according to manufacturer’s instructions (Abcam, ab102526).

**Western Blots**

Following a PBS washing step, infected cell monolayers were lysed using 100 µL of cold RIPA buffer (10 mM Tris-HCl, pH 9.0, 1 mM EDTA, 1% Triton X-100, 0.1% sodium deoxycholate, 1% SDS, 140 mM NaCl, prepared in Milli-Q water). Sample protein concentrations were quantified and normalized using a commercial colorimetric assay (Bio-Rad, Hercules, CA, USA; Cat. No. 5000111). The addition of 6× cracking buffer (0.35 M Tris pH 6.8, 30% glycerol, 10% SDS, 0.125% bromophenol blue, 0.05% 2-mercaptoethanol) to each sample, followed by a 10 min heat-block boiling step, was performed. Samples were then run on an SDS-PAGE gel consisting of a separating gel (10% acrylamide/bis solution, 29:1, 3.76 M Tris; 0.1% SDS; 0.1% APS; 0.1% TEMED) and a stacking gel solution (5.12% acrylamide/bis solution, 29:1, 0.13 M Tris; 0.1% SDS; 0.1% APS; 0.1% TEMED), while nitrocellulose membrane transfer, blocking, and antibody incubations (antibodies listed in Table 2) were performed, according to previously described protocols [9].

| Target | Host | Conjugate | Company | Catalog |
| --- | --- | --- | --- | --- |
| p-TBK-1 | Rabbit IgG | Unconjugated | Cell Signaling Technology, Danvers, MA, USA | 5483S |
| TBK-1 | Rabbit IgG | Unconjugated | Cell Signaling Technology, Danvers, MA, USA | 3504S |
| pIRF3 | Rabbit IgG | Unconjugated | Invitrogen, Carlsbad, CA, USA | PA5-36775 |
| IRF3 | Rabbit IgG | Unconjugated | Cell Signaling Technology, Danvers, MA, USA | 4302S |
| pIRF7 | Rabbit IgG | Unconjugated | Cell Signaling Technology, Danvers, MA, USA | 5184S |
| IRF7 | Rabbit IgG | Unconjugated | Cell Signaling Technology, Danvers, MA, USA | 4920S |
| pSTAT-1 | Rabbit IgG | Unconjugated | Cell Signaling Technology, Danvers, MA, USA | 9167S |
| pSTAT-2 | Rabbit IgG | Unconjugated | Cell Signaling Technology, Danvers, MA, USA | 88410S |
| MDA-5 | Rabbit IgG | Unconjugated | Cell Signaling Technology, Danvers, MA, USA | 5321S |
| RIG-I | Rabbit IgG | Unconjugated | Cell Signaling Technology, Danvers, MA, USA | 3743S |
| OAS-1 | Rabbit IgG | Unconjugated | Cell Signaling Technology, Danvers, MA, USA | 14498S |
| B-actin | Mouse IgG | Unconjugated | Millipore Sigma, Burlington, MA, USA | MAB1501 |
| Rabbit IgG [H] + [L] chains | Goat IgG | HRP | Cell Signaling Technology, Danvers, MA, USA | 7074S |
| Mouse IgG [H] + [L] chains | Horse IgG | HRP | Cell Signaling Technology, Danvers, MA, USA | 7076S |

**Supplementary Table 2:** List of antibodies used in western blot experiments.

**Immunoassays**

Levels of cytokines and chemokines were measured by a combination of multiplex immunoassays and single-analyte ELISAs. Commercial ELISA kits were used to analyze the expression levels of IFNα (Invitrogen, BMS216), IFNβ (PBL Assay Science, 41410), TNFa (Invitrogen, KAC1751) and IL-12 p70 (Invitrogen, BMS238). For cell culture supernatants, the V-PLEX Human Proinflammatory Panel 1 (Meso Scale Discovery, K15049D) and V-PLEX Human Cytokine Panel 1 (Meso Scale Discovery, K15050) were used to quantify the levels of proinflammatory cytokines and chemokines according to the manufacturer’s instructions, on a SQ 120 MSD instrument. For the measurement of analytes in macaque plasma and BAL, the ProcartaPlex Non-Human Primate Cytokine/Chemokine/Growth Factor Panel 37plex (Invitrogen, EPX370-40045-901) was used on longitudinal samples from all 6 macaques and read the assay on an xMAP Intelliflex System (Luminex). Standard curve analysis was performed using the ProcartaPlex™  Analysis App on Thermo Fisher Connect. All samples were assayed in duplicate, and serially diluted standards were included in each assay.

Antibodies used in cell culture include anti-IFNAR2 (Invitrogen, 213851), anti-IFNGR1 (Invitrogen, PA5-47866), anti-CD46 (Millipore-Sigma, MABF2182).

**Electron Microscopy**

For ultrastructural observations of infected cells by thin sections, samples were fixed in 2.5% glutaraldehyde in 0.1 mM sodium cacodylate and processed as described previously [10]. Ultrathin sections of infected cells were stained with osmium tetraoxide before examination with Hitachi 7600 EM under 80 kV equipped with a dual AMT CCD camera system. Quantitative measurement of length for the inclusion membrane and host ER elements attached to this membrane using ImageJ was performed on 24 representative electron micrographs at low magnification to ensure the entire inclusion fit into the field of view.

**Reverse Genetics**

Recombinant viruses were generated from previously described protocols [11]. Briefly, gene fragments from IDT were assembled into vector plasmids, pMV^EZ^ or pCG^EZ^ that were then ligated together after PCR amplification of homologous regions to generate a full-length rMV^EZ^ antigenomic plasmid. Recombinant viruses were rescued from MVA-T7-infected Vero-SLAM cells transfected with full-length plasmid along with plasmids expressing MeV proteins N, P and L. Recombinant viruses were tested for Mycoplasma contamination, mutated regions were tested by Sanger sequencing, and viral titers were measured by plaque assay.

**Immunofluorescent Microscopy**

Infected PBMCs were washed and resuspended in 1x PBS, then overlayed on coverslips for 1 hour. Next, the cells were fixed in 4% paraformaldehyde for 10 min at RT. After fixation, the cells were washed with PBS and permeabilized with PBS containing 0.3% Triton X-100 for 10 min. The cells were blocked with permeabilization buffer supplemented with 3% BSA for 30 min, and were then incubated with primary antibody (anti-MeV N-FITC – see Table 3) in blocking buffer for 1 h, washed, and resuspended in ProLong Gold Antifade reagent with DAPI (Invitrogen, P36931). The cells were examined using a Leica Thunder imaging system, at 60x magnification. Images were analyzed using the Leica Thunder Analysis Suite and Image J software for counting.

**Flow Cytometry**

Cells were harvested at 24 hours post-infection, washed in PBS and resuspended in FC-block (Invitrogen, 14-9161-73) with Live-Dead stain (Biolegend, 423105) for 10 minutes. After more PBS washes, the cells were stained with extracellular antibodies for 20 minutes, followed by fixation and permeabilization (Invitrogen, 00-5523-00) and then stained with intracellular antibodies for 20 minutes. Finally, cells were resuspended in 1% PFA and read on a Cytek Northern Lights flow cytometer (Cytek). Antibodies are listed in Table 3.

| Target | Clone | Fluorophore | Company | Catalog Number |
| --- | --- | --- | --- | --- |
| CD4 | L200 | BV421 | BD BioSciences | 562842 |
| CD123 | 7G3 | AF405 | BD BioSciences | 560087 |
| CD66abce | TET2 | VioBlue | Miltenyi Biotec | 130-119-938 |
| CD11c | 3.9 | BV480 | BD BioSciences | 565806 |
| CD11b | ICRF44 | BV570 | BioLegend | 301325 |
| CD14 | M5E2 | BV650 | BD BioSciences | 563419 |
| CD20 | 2H7 | BV711 | BD BioSciences | 563126 |
| CD150 | A12 | BV786 | BD BioSciences | 744348 |
| MeV N | 83KKII | FITC | Millipore Sigma | MAB8906 |
| CD8a | SK1 | Spark Blue 550 | BioLegend | 344759 |
| CD38 | OKT10 | PE | BD BioSciences | 367146 |
| CCR7 | G043H7 | PE-Dazzle594 | BioLegend | 353235 |
| HLA-DR | L243 | PE-Fire640 | BioLegend | 307675 |
| CD16 | 3G8 | BB700 | BD BioSciences | 746199 |
| FoxP3 | 259D/C7 | RB705 | BD BioSciences | 570239 |
| CD46 | 1C6 | PE-Cy7 | BD BioSciences | 665118 |
| CD138 | DL-101 | AF647 | BioLegend | 352313 |
| CD27 | O323 | Spark NIR 685 | BioLegend | 302855 |
| CD159a | REA110 | Vio Bright R720 | Miltenyi Biotec | 130-128-577 |
| CD3 | SP34-2 | BUV737 | BD BioSciences | 568353 |
| Live/Dead |  | Zombie NIR | BioLegend | 423105 |
| CD45RA | 5H9 | APC-H7 | BD BioSciences | 561212 |

**Supplementary Table 3:** List of antibodies used for flow cytometry.

**Statistical Analysis**

For statistical analyses, two-tailed Student’s t test (α = 0.05) was used for nonparametric normally distributed data while analysis with Mann-Whitney U test was used for nonparametric data to compare two groups. Analysis of variance (ANOVA) with Bonferroni post hoc tests was used for comparing multiple groups. All data were analyzed using GraphPad Prism software. Statistical significance was determined as *P < 0.05, **P < 0.01, and ***P < 0.001.

**Supplemental Figures:**

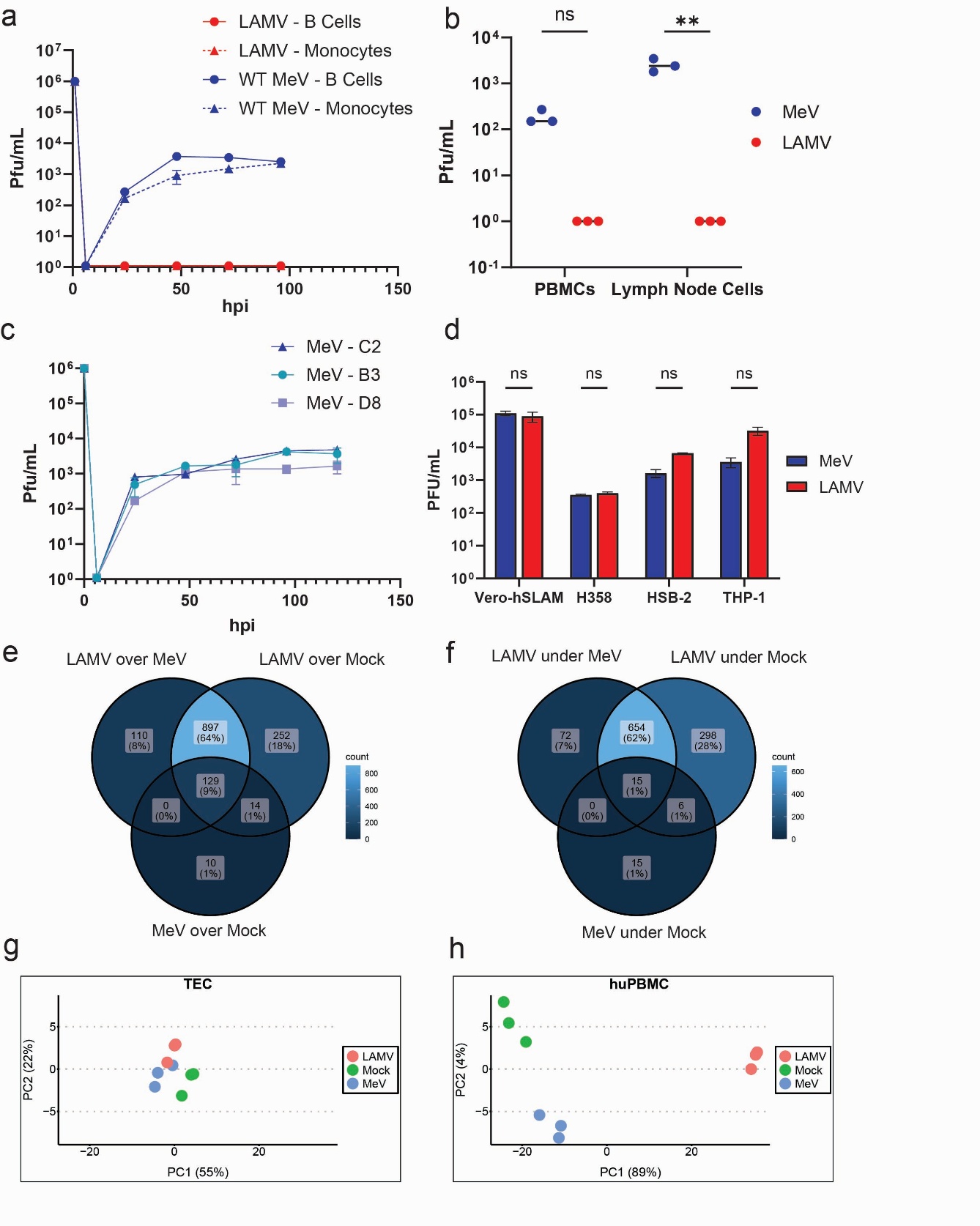
**Supplemental Figure 1. (a)** Infectious viral titers from the supernatant of magnetically-isolated B cells and monocytes infected with either MeV or LAMV (MOI=1). **(b** Infectious virus titers, 72hours post-infection with MeV or LAMV, from the supernatant of lymph node cells and PBMCs derived from 3 different uninfected rhesus macaques. **(c)** Infectious virus titers for 3 different strains of MeV, including currently circulating strains (B3 and D8 genotypes). **(d)** Infectious virus titers, 72 hours post infection with MeV or LAMV, from the supernatant of four different immortalized cell models. **(e-f)** Venn diagrams of over-expressed **(e)** and under-expressed **(f)** genes across all three conditions in huPBMC infection, with log2FC > 1 and pval > 0.01. **(g-h)** PCA plots of normalized transcript counts during infection of TECs **(g)** and huPBMCs **(h)** with either MeV, LAMV or mock-infection.

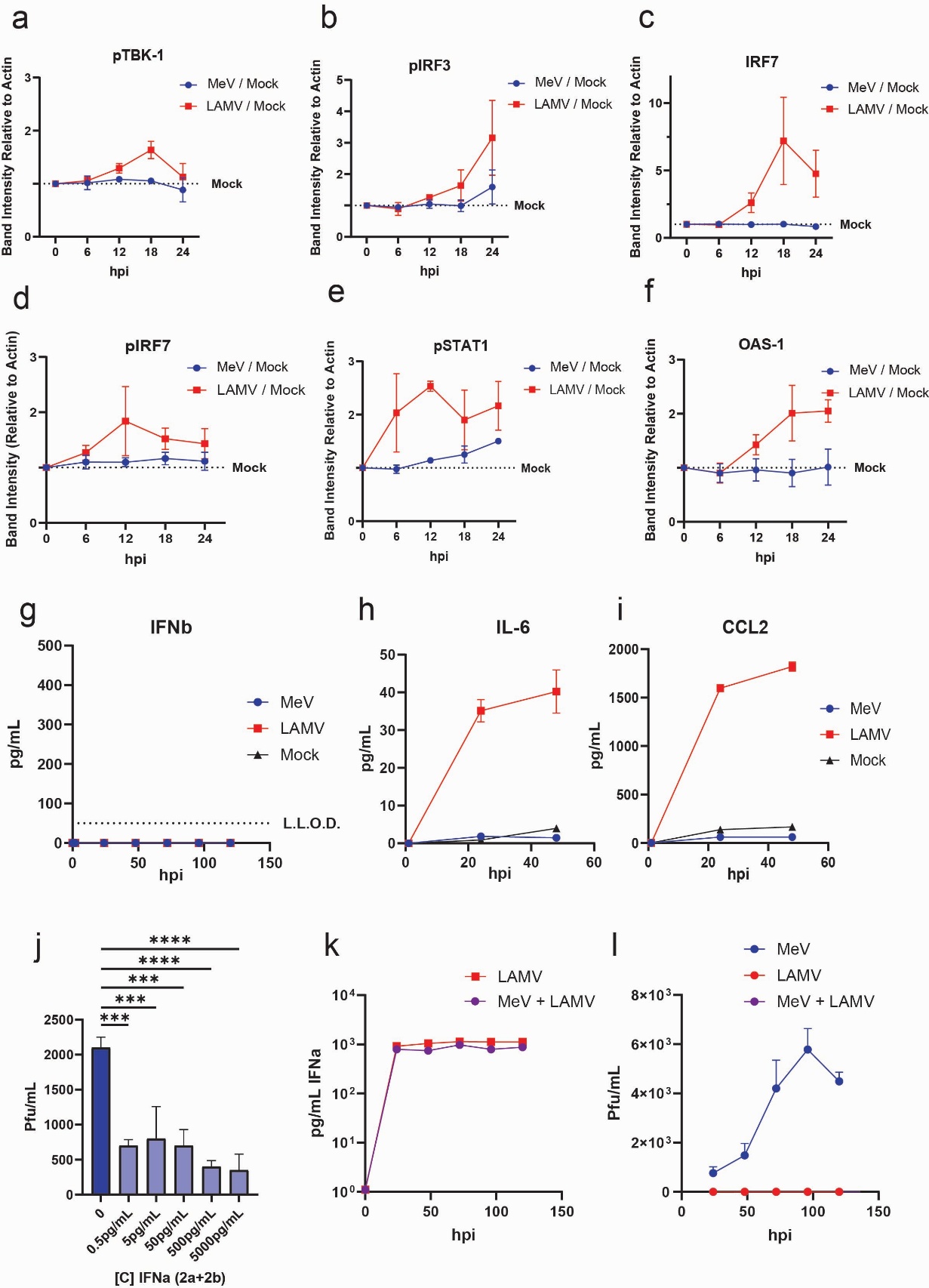

**Supplemental Figure 2. (a-f)** Densitometry analysis of western blotting for phosphorylated TBK-1 **(a)** phosphorylated IRF3 **(b)**, IRF7 **(c)**, phosphorylated IRF7 **(d)**, phosphorylated STAT-1 **(e)**, OAS-1 **(f)** - normalizing pixelation to mock-infected cells and actin loading controls. The dotted line represents baseline mock levels. **(g)** IFNβ secretion during huPBMC infection, by ELISA. **(h-i)** IL-6 **(h)** and **(i)** CCL2 secretion during huPBMC infection, by multiplexed immunoassay. **(j)** IFNα secretion during infection of huPBMCs with LAMV or LAMV and MeV together (MOI=1). **(k)** Infectious virus titers during infection of huPBMCs with LAMV or LAMV and MeV together (MOI=1). **(l)** Infectious MeV titers, 72 hours post-infection with titration of IFNα subtypes 2a and 2b at different concentrations.

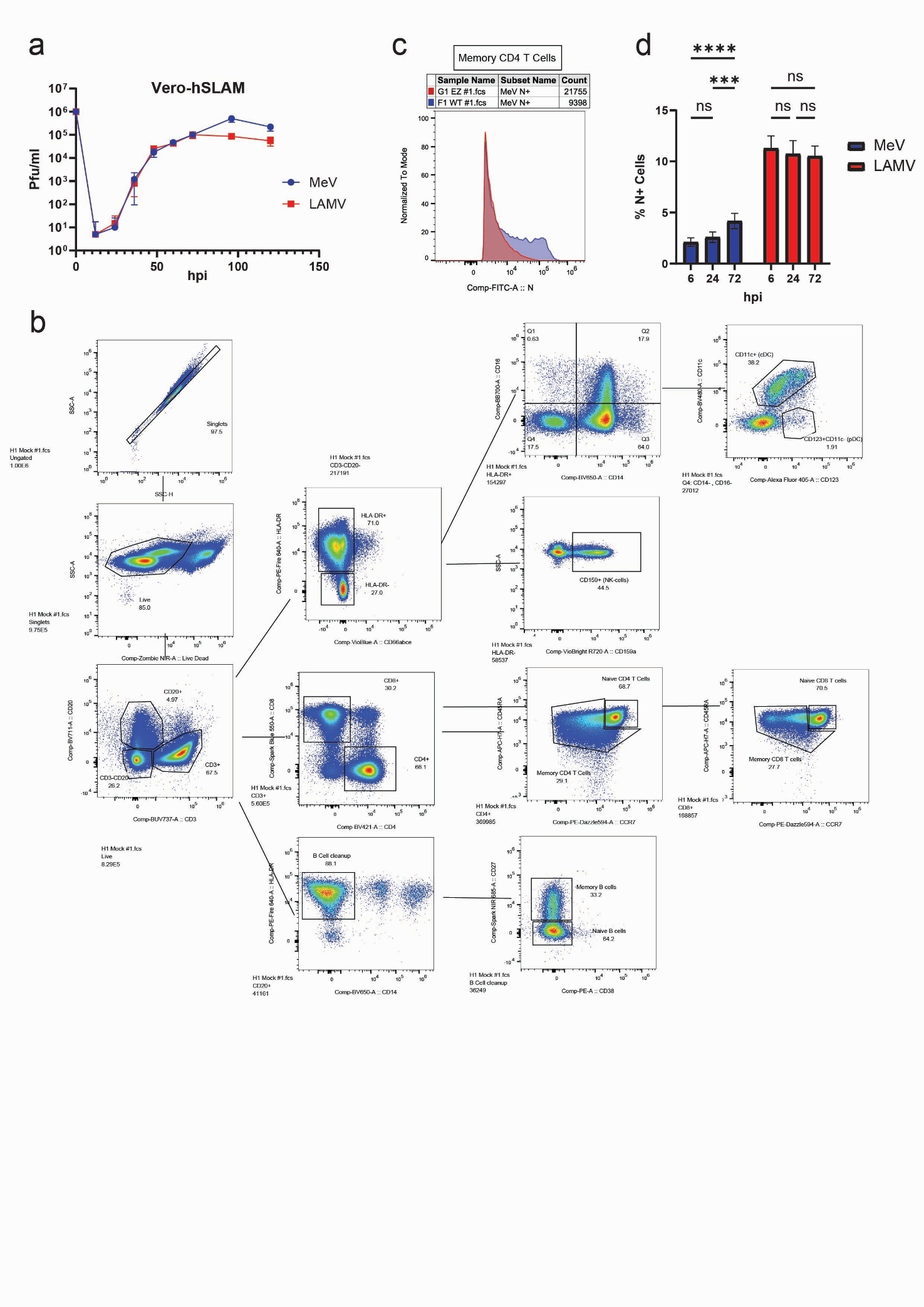

**Supplemental Figure 3. (a)** Infectious virus titers in vero-hSLAM cells infected with MeV or LAMV (MOI=0.01). **(b)** Overlay of FITC (N) MFI histograms for memory CD4 T cells, infected with either MeV (blue) or LAMV (red). **(c)** Percentage N+ huPBMCs infected with MeV, LAMV or mock infection, by immunofluorescent staining at 6, 24, and 72 hours post-infection. **(d)** Gating scheme for subset determination during flow cytometry of huPBMCs.

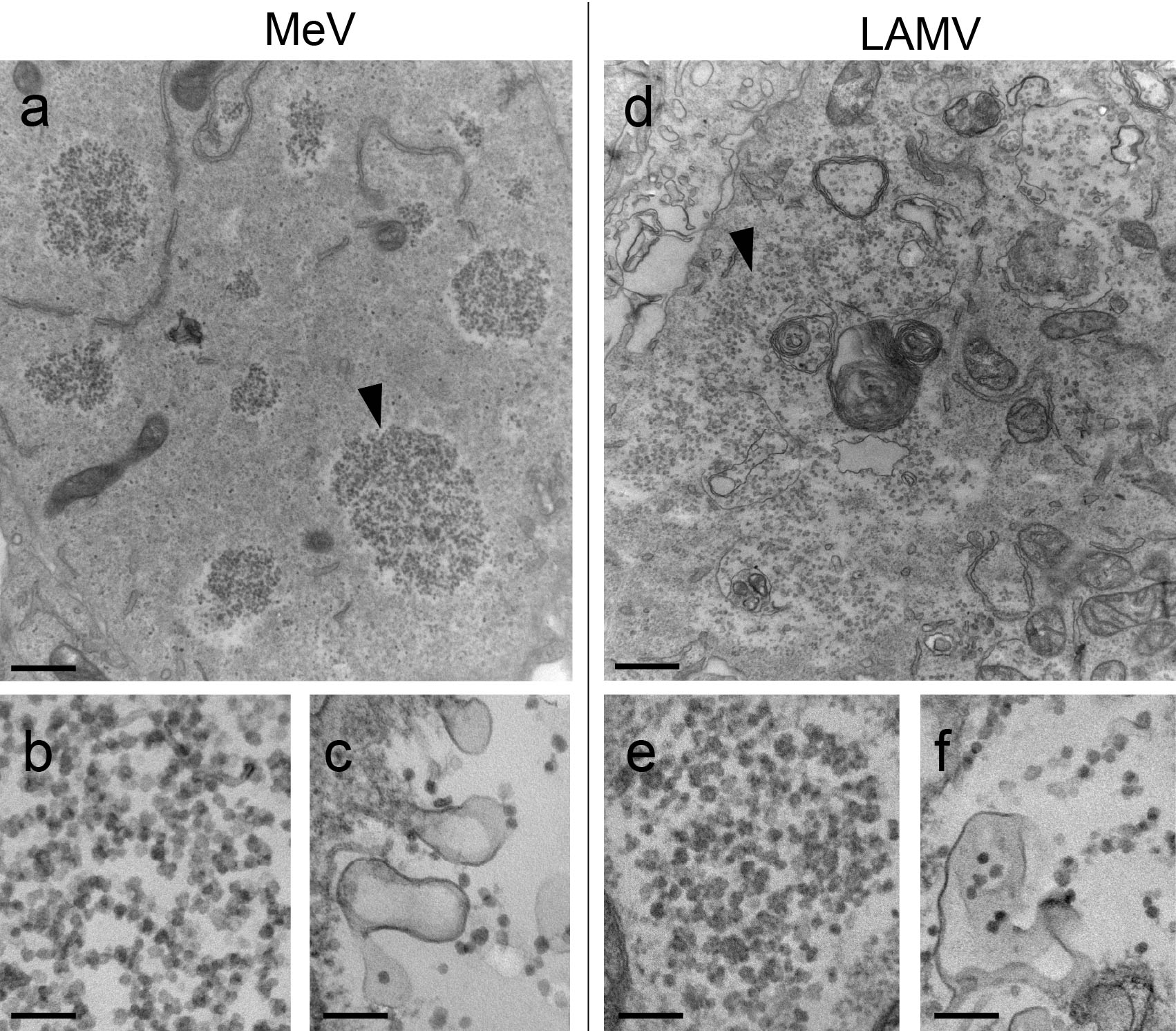

**Supplemental Figure 4. (a-f)** Ultrastructure of LAMV or MeV in vero-hSLAM cells. Cells were infected for 60h for EM observations showing at low magnification many sites of viral replication (arrowheads) of MeV **(a)** or LAMV **(d).** Higher magnification of viral RNPs for MeV **(b)** and LAMV **(e)** inside cells**.** Egressing viruses were observed for both strains in the extracellular medium **(c and f)**. Scale bars, 0.1 µm, except 0.5 µm for a and d.

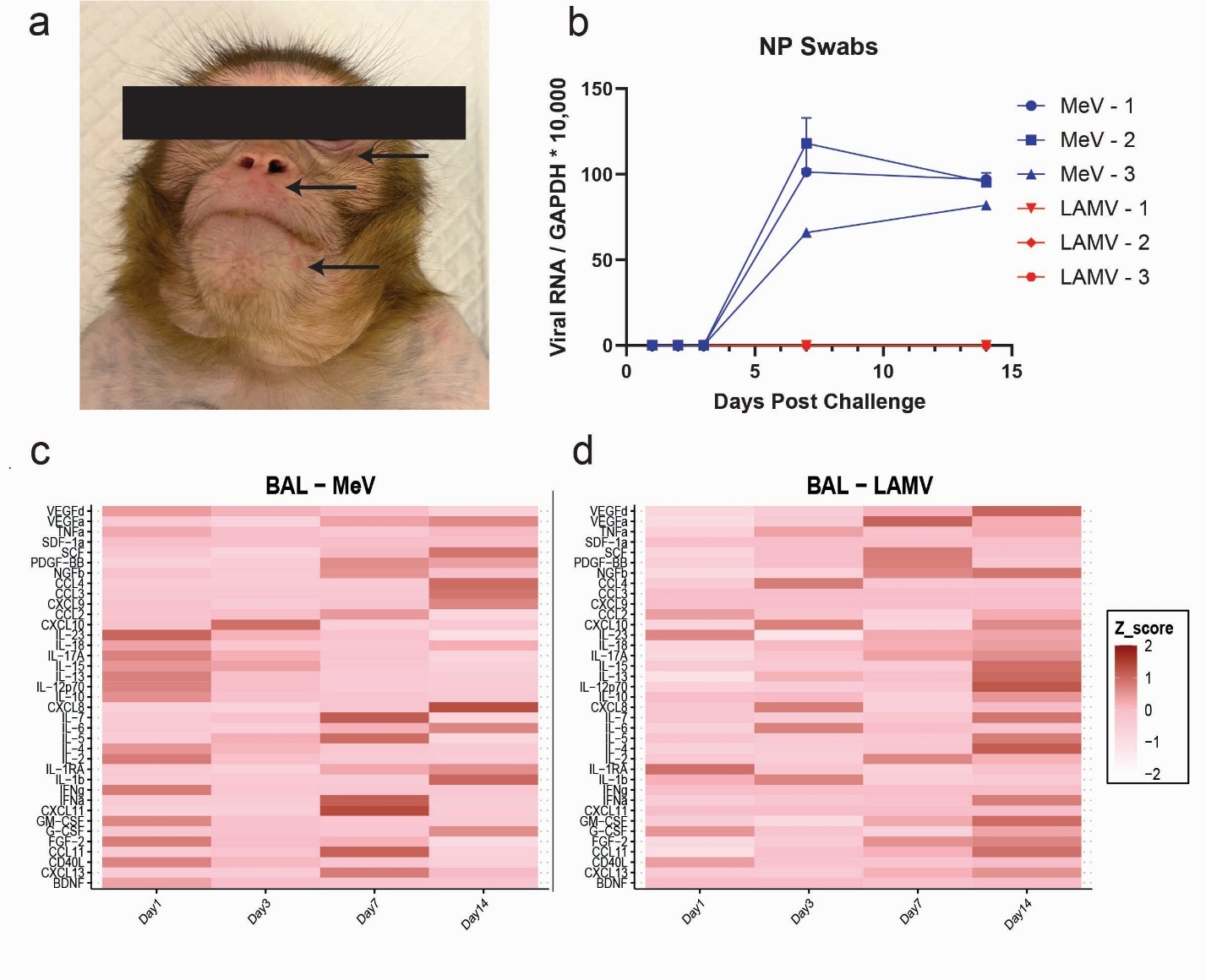

**Supplemental Figure 5. (a)** Example of Measles rash (arrows), 11 days post-challenge on a MeV-infected macaque. **(b)** RTqPCR of viral RNA from cells collected in nasopharyngeal swabs for all 6 rhesus macaques. Normalized to GAPDH levels. **(c-d)** Heatmap of z-scores of cytokine levels in BAL plasma of both MeV **(c)** and LAMV **(d)** infected macaques, normalized to each macaque’s own baseline expression.

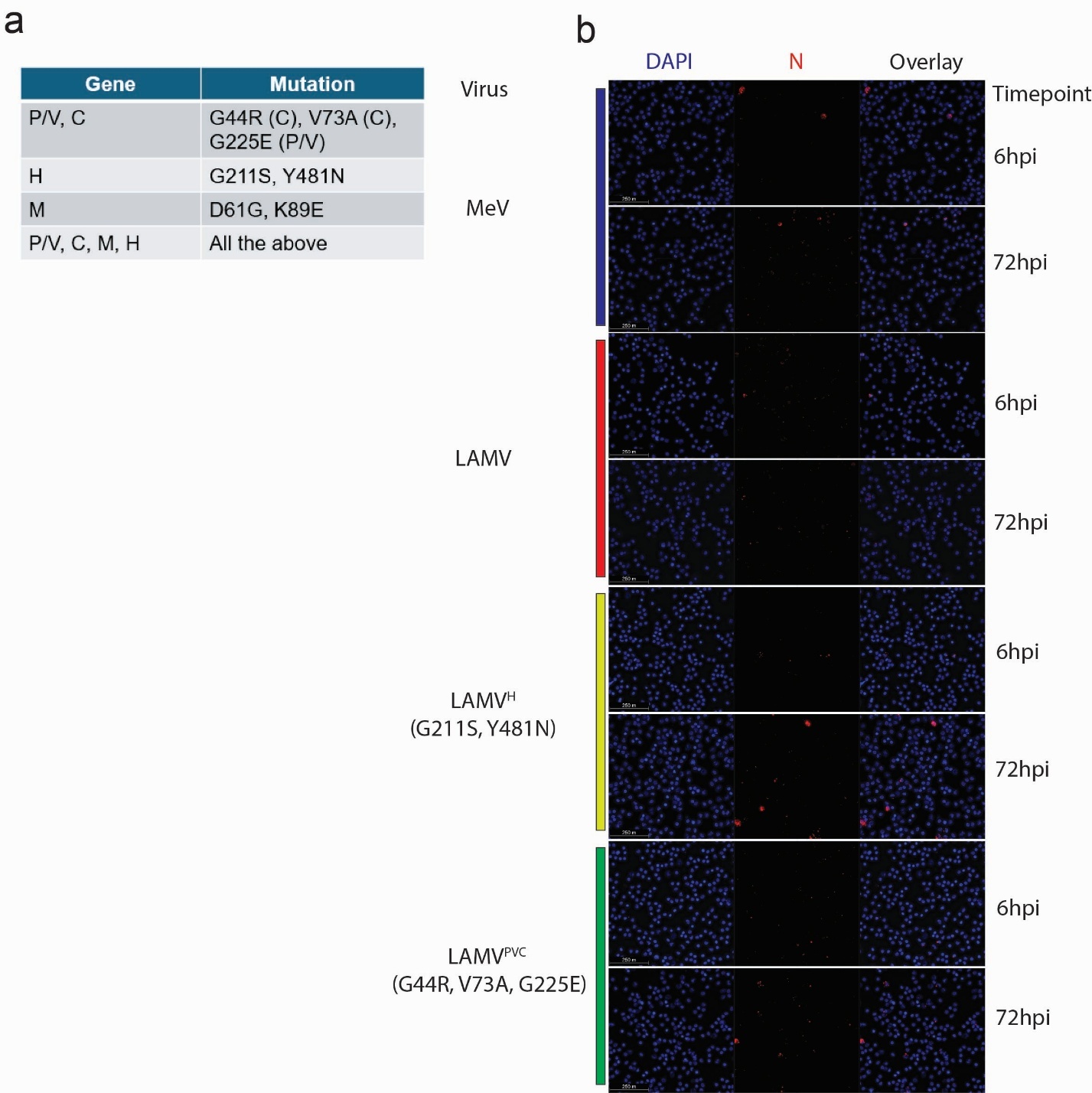

**Supplemental Figure 6. (a)** Table of conserved LAMV mutations that were reverted as WT amino acids into the LAMV backbone. **(b)** IF staining for viral protein N in huPBMCs during infection with different viruses at 6 and 72 hours post-infection (MOI=1), with nucleus staining in blue and N staining in red.
